# Memory drives the formation of animal home ranges: evidence from a reintroduction

**DOI:** 10.1101/2020.07.30.229880

**Authors:** Nathan Ranc, Francesca Cagnacci, Paul R. Moorcroft

## Abstract

Most animals live in a characteristic home range, a space-use pattern thought to emerge from the benefits of memory-based movements; however, a general model for characterizing and predicting their formation in the absence of territoriality has been lacking. Here, we use a mechanistic movement model to quantify the role of memory in the movements of a large mammal reintroduced into a novel environment, and to predict observed patterns of home range emergence. We show that an interplay between memory and resource preferences is the primary process influencing the movements of reintroduced roe deer (*Capreolus capreolus*). Our memory-based model fitted with empirical data successfully predicts the formation of home ranges, as well as emerging properties of movement and revisits observed in the reintroduced animals. These results provide a quantitative framework for combining memory-based movements, resource preference and the emergence of home ranges in nature.

## Introduction

Most animals live in home ranges – areas that are typically much smaller than their movement capabilities would otherwise allow^1^. The spatially-constrained nature of animal space-use has important implications for many ecological processes, including density-dependent regulation of population abundance^2^, predator-prey dynamics^3^, the spread of infectious diseases^4^, as well as for the design of conservation strategies^5^. Home ranges are pervasive throughout the animal kingdom, suggesting that they may provide fitness benefits in a wide range of ecological contexts, and originate from general biological mechanisms^6^. In territorial species, the emergence of a constrained space-use has been successfully characterized by analytical movement models based on conspecific avoidance^3,7–9^. However, a general model for predicting emergent patterns of space-use is still lacking for animals that form home ranges in the absence of territoriality or central place foraging.

In recent years, increasing attention has been devoted to the hypothesis suggesting that home ranges emerge from the foraging benefits of memory^10,11^. Theoretical studies have demonstrated the foraging advantages of memory over proximal mechanisms (e.g., area-restricted search and perception) in spatially-heterogeneous, predictable landscapes^12–14^. In turn, simulations have shown that memory-based movements can lead to the formation of stable home ranges^15,16^, and to non-territorial spatial segregation between individuals^17^. However, our understanding of how memory influences animal movement and resulting space-use patterns in nature is still in its infancy.

Optimal foraging experiments have provided evidence for the adaptive value of memory. For example, green-backed fire-crown hummingbirds (*Sephanoides sephaniodes*) can achieve substantial energy gains by adjusting their visit frequency to the renewal dynamics of high-quality resources at memorized locations^18^. At larger spatial scales, mechanistic models based on telemetry data have shown that animals are capable of memorizing the location and profitability of resources^19,20^. For example, roe deer (*Capreolus capreolus*) rely on memory, and not perception, to track the dynamics of resource availability within their home range^21^. Whether mechanistic movement models parametrized with empirical data can capture the spatial patterns of animal home ranges in nature remains, however, largely unanswered.

Most studies of animal home range movements have been conducted on resident animals whose experience and knowledge of the surrounding environment is already well-developed at the onset of monitoring^19,20,22,23^. This is problematic when studying the effects of memory because animals are utilising knowledge obtained prior to the observation period, which has been proposed as the reason for discrepancies between memory-based movement model predictions and observed space-use patterns^22^. One approach to address this challenge is to examine the process of home range formation (also referred to as emergence) when animals have been introduced into a novel environment^11^, where it can be reasonably assumed that the animals have no existing memories of the local environment.

In this study, we elucidate the role of memory in the movements of animals by analysing the process of home range formation of individuals reintroduced into a novel environment. Our results show how the interplay between memory and resource (landscape attributes) preferences gives rise to observed patterns of home ranges. Specifically, we fit an individual-based, spatially explicit movement model to the observed trajectories of European roe deer reintroduced into the Aspromonte National Park (Calabria, Italy), where the species had previously been extirpated. This experimental system is ideally-suited to the study of the biological determinants of home ranging behaviour for three reasons. First, because roe deer were released into a novel environment, as noted above, the theoretical challenge of how to initialize memory at the beginning of the simulation is essentially side-stepped. Second, because roe deer are solitary^24^, their movements are expected to be primarily based on individual information rather than group decision making^25^. Third, because roe deer population was being re-established, animal density was low throughout the study, therefore limiting the influence of intraspecific competition on individual movements and space-use.

Roe deer were fitted with GPS telemetry collars and monitored from their release into the study area till the collars ceased functioning (n =17 individuals; see *Methods*). We analysed the biological processes underlying roe deer movements (n = 17,136 six-hour movement steps) using a redistribution kernel^9,20,26^. The model characterizes the probability that a given individual moves from its current position to any location in the landscape as a function of motion capacity, and a weighting function including resource preferences and memory. Building up on earlier work^15–17^, memory was represented as a bi-component mechanism: a *reference memory* encoding long-term attraction to previously visited locations, and a *working memory*, which accounts for a short-term avoidance of recently visited locations (for example, due to local resource depletion^15^). The dynamics of both memory components are governed by their respective learning and decay rates, and associated spatial scale.

We hypothesized that the interplay between memory and resources was the primary driver underlying roe deer movements (H1). To this end, we fitted two competing movement models: (i) a *resource-only* model (M_res_) in which roe deer movement was only influenced by resource preferences (which in this case corresponds to landscape attributes such as slope, tree cover and landcover categories; sensu^27^). (ii) a *memory-based* model (M_mem:res_) in which movement was governed by the interplay between memory and resource preference (sensu^26^). Following on our previous work that examined memory dynamics in an experimental setting^21,28^, we predicted that the empirical movement data would provide a higher support to the memory-based model than its resource-only counterpart (P1.1). In addition, we predicted that, overall, roe deer would strongly select for previously visited locations (P1.2).

We further hypothesized that the interplay between memory and resource preferences can lead to formation of home ranges, as observed in the reintroduced roe deer (Cagnacci *et al*. in prep; H2). To this end, we compared the emerging movement and space-use properties of trajectories simulated from the parametrized redistribution kernels with those from the empirical roe deer movements. Accordingly, we predicted that, in contrast with the resource-only model, the simulations from the memory-based model would lead to spatially-constrained movements (P2.1) with a high prevalence of acute turning angles (P2.2). In addition, we predicted that memory-based movements would be characterized by a high number of revisitations (also referred to as movement recursions^29^; P2.3). Further details on the mathematical formulations of the redistribution kernel and on the movement simulations can be found in the *Methods* section.

## Results

### Biological drivers of reintroduced roe deer movements

The movement model that included both memory and resource preferences (M_mem:res_) had overwhelmingly stronger support compared to the resource-only model (M_res_; Δlog-likelihood = – 8684; Δdf = −6; ΔAIC = 17355; p-value < 0.001; Table 1; P1.1 supported). Memory was a key biological process underlying the movements of reintroduced roe deer (most influential variable; Table 1; P1.1 supported). The importance of memory was primarily due to the effects of reference memory (ΔAIC = 1278 when working memory was removed, compared to ΔAIC = 17355 when both working and reference memory were removed; see Table 1). With respect to reference memory, the spatial scale of learning was most influential (ΔAIC = 7444 if learning occurred only on the visited locations i.e., *λ_R_* = ∞), followed by learning rate (ΔAIC = 3434 if learning was immediate i.e., *l_R_* = 1) and decay rate (ΔAIC = 2631 if there was no memory decay i.e., *δ_R_* = 0).

**Table 1.**
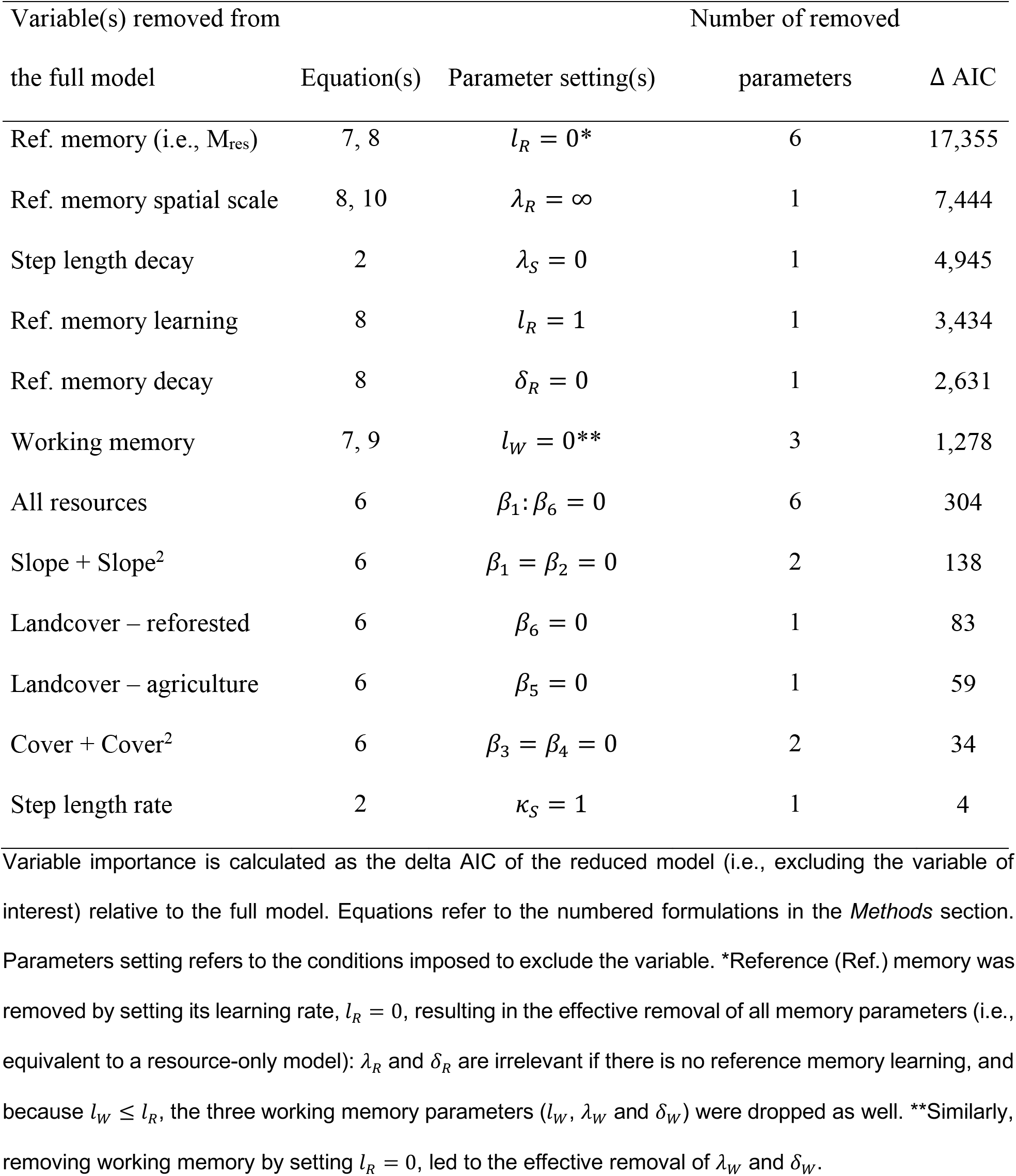
Variable contributions to the memory-based model (M_mem:res_).

Roe deer acquired memories of visited locations, with the learning curve reaching half its maximum value (i.e., memory = 0.5) after 7.9 days for reference memory (*l_R_* = 0.0217 6h^-1^; see Fig. 1a for confidence intervals), and after 8.4 days for working memory (*l_w_* = 0.0204 6h^-1^). Spatially, information was gained beyond the visited locations: reference memory learning decayed with distance to half its maximum value at 14.3 m (2# = 0.0485 m^-1^; meaning that at 25 m distance, learning was approximately 30 % that of the amount of memory acquired on the visited spatial location). Working memory learning declined to half its maximum value at 8.1 m (*λ_w_* = 0.0855 m^-1^; meaning that the learning rate at 25 m distance was approximately 12 % that of the visited location). Temporally, reference memory decayed with time since last visit with a half-life (t_1/2_) of 9.5 days (*δ_R_* = 0.0182 6h^-1^) while working memory decay was nearly instantaneous (t_1/2_ < 1 h; *δ_w_* = 0.99 6h^-1^).

**Figure 1:**
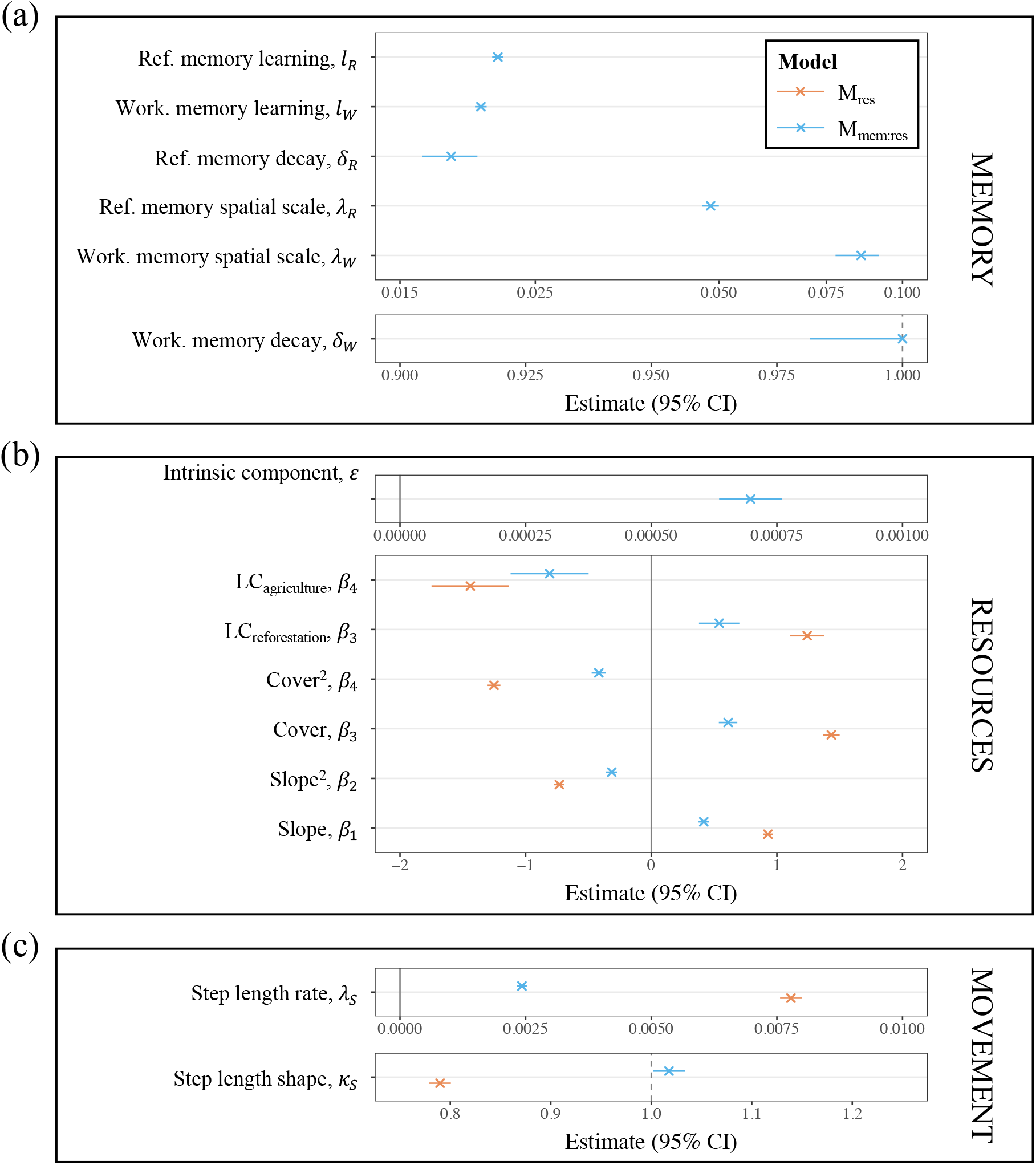
Parameter estimates. The estimates for the resource-only (M_res_; orange), and the memory-based (M_mem:res_; blue) models are plotted with the corresponding 95% marginal confidence intervals. Memory (panel a), resource preference (b) and movement (c) parameters are shown separately for readability.

The combined effect of memory dynamics and of the intrinsic component of resource preference (i.e., the attraction of locations in absence of memory; *ε* = 6.94×10^-4^; Fig. 1b) led to a very strong selection for previously-visited locations (P1.2 supported). Specifically, the first visit of a given location resulted in a 31.6-fold increase in its attraction, and a 10.1-fold increase on the adjacent locations (Fig. 2a).

**Figure 2:**
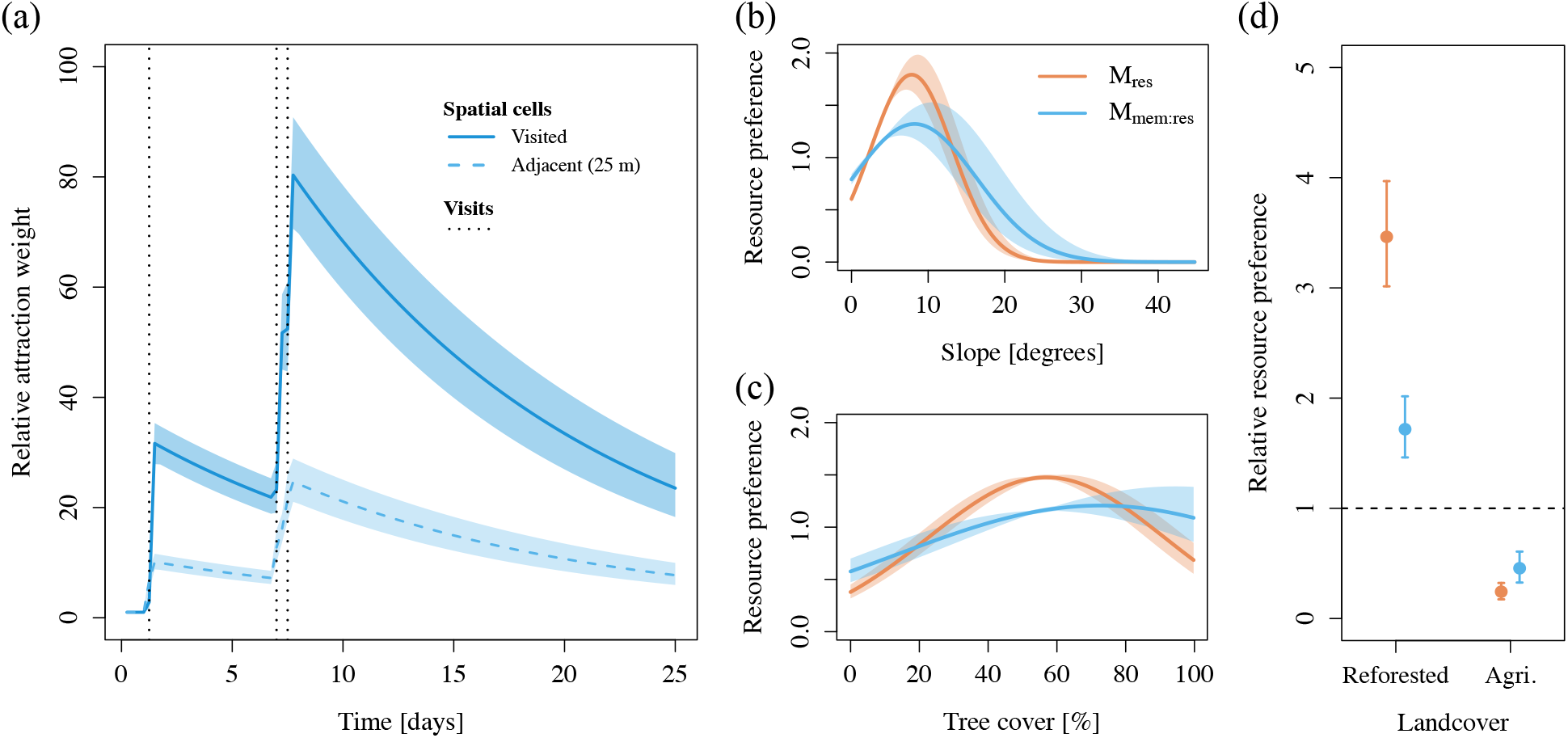
Predictor effects. The response curves for the resource-only (M_res_; orange) and the memory-based (M_mem:res_; blue) models are plotted with the corresponding 95% marginal confidence intervals. Panel (a) shows the attraction of a visited spatial cell (continuous line) and an adjacent cell (25 m away; dashed line) relative to a cell that has never been visited (attraction = 1) resulting from the fitted memory-based model. Hypothetical visits (at t = 1.25, 7.00 and 7.50 days) are shown in dotted vertical lines. Panel (b) and (c) illustrate the preference for slope and tree cover, respectively. Panel (d) shows the relative preference for reforested and agriculture landcovers.

Roe deer movements were also influenced by their resource preferences (Fig. 1b; ΔAIC = 304 when resource preferences were null i.e., *β*_1_:*β*_0_ = 0; Table 1). Roe deer preferred intermediate slopes (Fig. 2b), with peak preference at eight degrees (most influential resource; ΔAIC = 138 when *β*_3_ = *β*_2_ = 0; Table 1). Roe deer preference for tree cover was characterized by slight qualitative differences between models: preference for intermediate tree cover (a clear peak at 57%) in the resource-only model, and for intermediate and high levels of tree cover (a broad peak at 73%) in the memory-based model (Fig. 2c; ΔAIC = 34 when *β*_3_ = *β*_4_ = 0; Table 1). In addition, roe deer strongly preferred reforested areas and avoided agricultural areas (Fig. 3c; ΔAIC = 83 when *β*_6_ = 0 and ΔAIC = 59 when *β*_5_ = 0, respectively; Table 1). For all evaluated resources, preferences had a lower effect size for the memory-based model than for the resource-only model (Fig. 1b).

**Figure 3:**
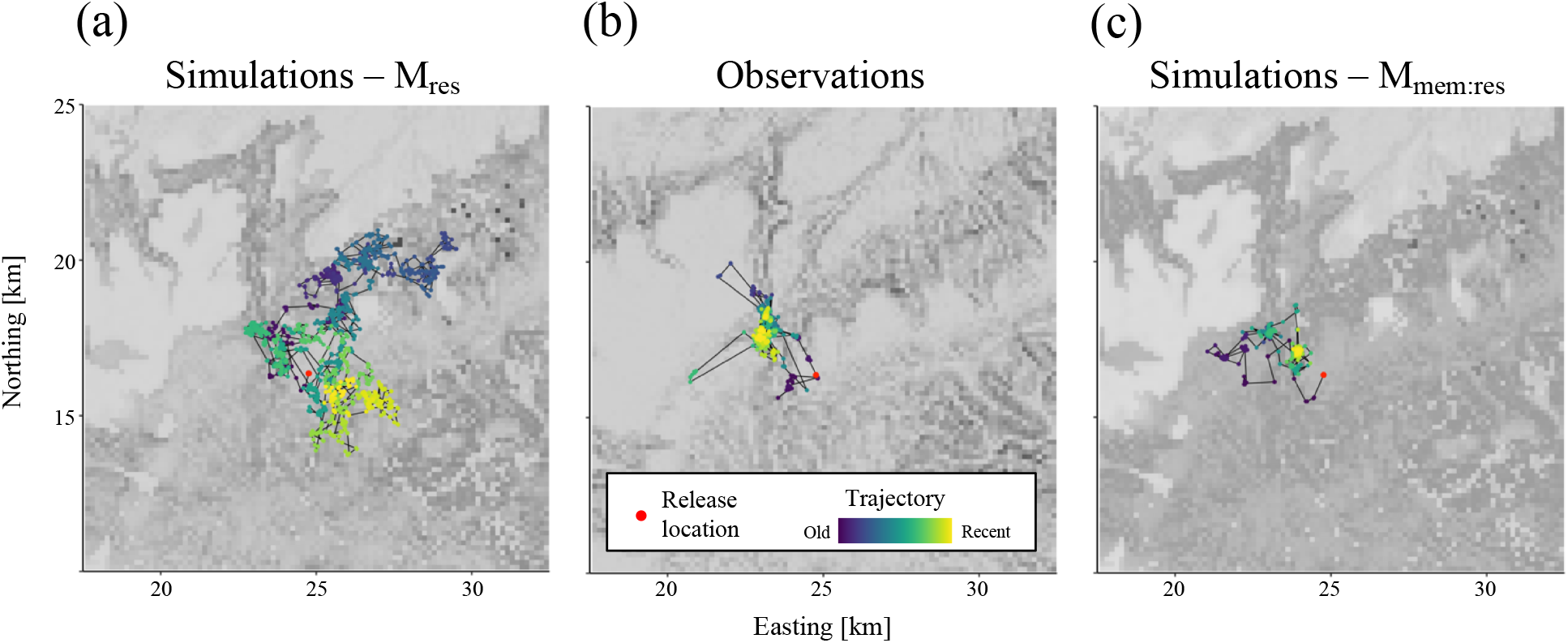
Movement trajectories. Three typical trajectories are shown for the resource-only simulations (M_res_; panel a), observed roe deer movements (panel b), and memory-based simulations (M_mem:res_; panel c). The release location is shown as a red dot and the time since release illustrated as a colour gradient (blue = old, yellow = recent). The trajectories were selected from the sample displayed on Figure 4.

Roe deer motion capacity greatly differed between the two competing movement models. The resource-only model characterized the movement distances between six-hour relocations as a heavy-tailed Weibull distribution (shape parameter *κ_S_* = 0.79; decay rate parameter *λ_S_* = 0.0078; Fig. 1c), with a corresponding mean step length of 147.0 m. In contrast, the memory-based model indicates a nearly 3-fold larger motion capacity (*κ_S_* = 1.02; *λ_S_* = 0.0024) corresponding to a mean step length = 409.4 m). The value of the shape parameter *κ_S_* being close to one implies that the step length distribution can be simplified to a negative exponential with a relatively small decrease in model accuracy (ΔAIC = 4 if *κ_S_* = 1.00; Table 1). Step length decay rate was, however, a highly influential parameter (ΔAIC = 4945 when compared with a resource-selection type movement kernel which assumes that roe deer spatial locations independently of their proximity i.e., *λ_S_* = 0; Table 1).

### Emergent space-use and movement properties

Most reintroduced roe deer settled into a constrained space (i.e., formation of a home range) as shown visually by the spatial concentration of their movements (Fig. 3b). The movement simulations from the resource-only model were typical of a random walk (technically, an inhomogeneous random walk due to the effects of resource preferences; Fig. 3a). In contrast, the memory-based model captured the characteristic space-use behaviour observed in released roe deer (Fig. 3c; see Supplementary S1 for additional movement trajectories).

The visual differences in patterns of movement behaviour seen in Figure 3 were characterized and quantified by examining the temporal trends in net squared displacement (NSD) with time since release (Fig. 4). The resource-only model did not capture the observed spatially-restricted movements of the released roe deer, with no saturation in the NSDs of individuals (Fig. 4a), and a linear increase of the mean NSD across individuals (compare the solid red line on Fig. 4b with the red line and grey shaded area in Fig. 4a). In contrast, the predictions of the memory-based movement model were consistent with the temporal trends in the observed movements of the released animals as demonstrated by the occurrence of prolonged plateaus in the NSD of individual animals (Fig. 4c; P2.1 supported), and the fact that the observed mean NSD across individuals is within the bounds of the predictions of the memory-based movement model (compare the solid red line on Fig. 4b with the grey area on Fig. 4c).

**Figure 4:**
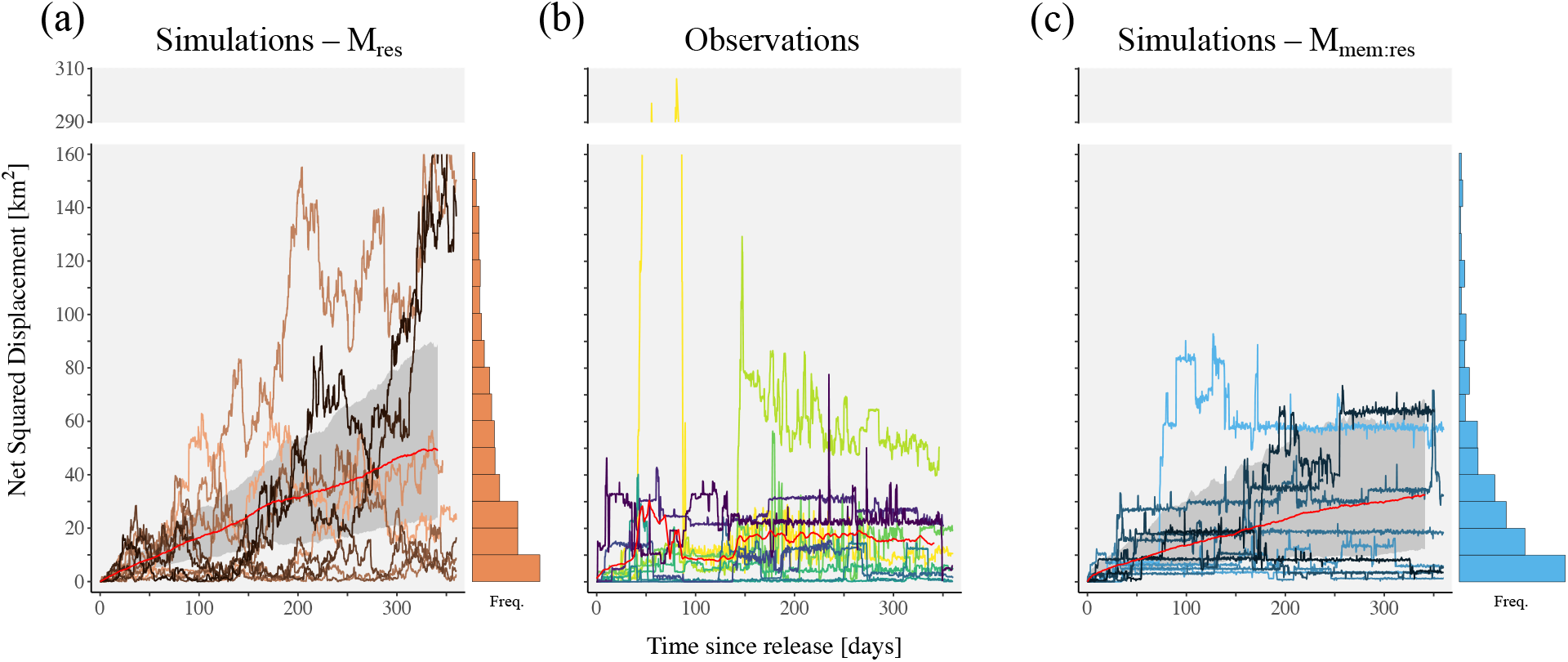
Trends in net squared displacement (NSD) with time since release. Panel (a): resource-only simulations (M_res_). Panel (b): observed roe deer movements. Panel (c): memory-based simulations (M_mem:res_). For the sake of clarity, only the individuals with more than 230 days of monitoring are shown (n = 10). For the simulations, one run for each of the selected individuals was randomly chosen. The trends in mean NSD across individuals are plotted as solid red lines (grey ribbons indicate the 5% and 95% bootstrapped quantiles for the simulations; panels a and c). The vertical histograms show the frequency of final NSD (i.e., evaluated at the end of the trajectories) for the simulations.

Both the resource-only and memory-based models had step length distributions that closely matched the observations (Fig. 5a,b). The memory-based model more accurately characterized the observed median step length while the resource-only model better captured its mean (median step lengths of 55.9, 75.0 and 50.0 m; means of 141.1, 135.2 and 105.5 m for observed movements, resource-only simulations and memory-based simulations, respectively). In contrast, the two models differed greatly in their ability to reproduce the observed patterns of turning angles: the resource-only model showed a uniform circular distribution of turning angles (Fig. 5c) whereas the memory-based model captured the high density of acute turning angles (in the vicinity of −*π* and +*π*), that are characteristic of observed roe deer movements (Fig. 5d; P2.2 supported).

**Figure 5:**
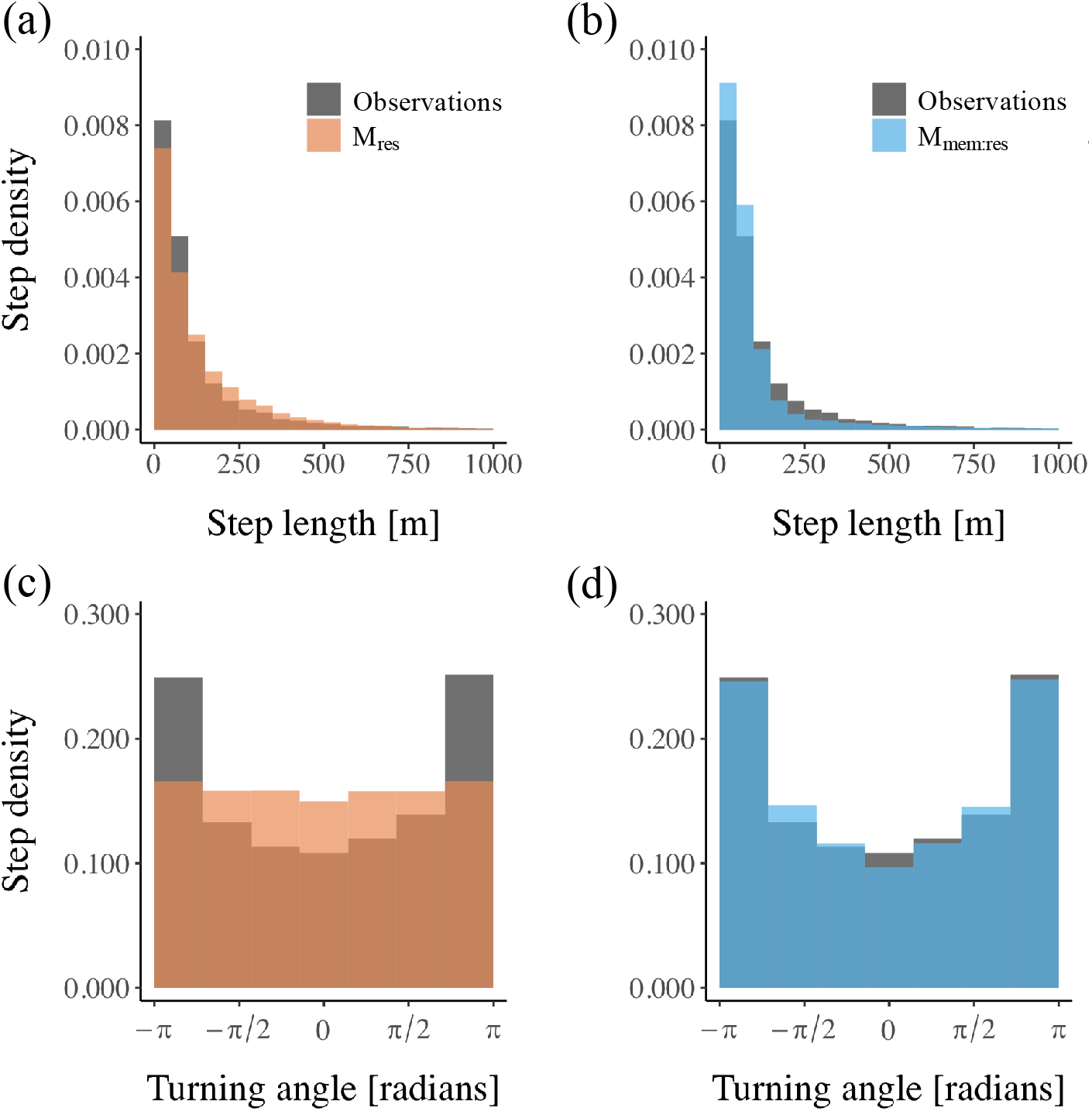
Emergent movement properties. The distributions of step length (panels a and b) and turning angle (panels c and d) are shown for observed roe deer movements (grey), the simulated trajectories from the resource-only model (M_res_; orange), and the simulated trajectories from the memory-based model (M_mem:res_; blue).

Observed roe deer movement behaviour was characterized by frequent revisits: 33.8% of the utilized locations (spatial scale = 25 x 25 m) were visited twice or more. The simulations from the resource-only model, however, had very few revisits – only 4.5% spatial locations were revisited – leading to a large mismatch with the revisitation patterns of observed trajectories (Fig. 6a). In contrast, the memory-based simulations were characterized by many revisits: 35.2% of the locations were revisited (P2.3 supported). The revisitation patterns of the memory-based model were highly similar to those of the observed roe deer movements (Fig. 6b), albeit with a slight tendency to underestimate the number of locations with few revisits (i.e., less than five), and to overestimate those with many revisits (especially above 20). Consistent with these patterns in the number of revisits, the memory-based model also captured the observed patterns of time since last visit more accurately than the resource-only model (compare Figs 6c and 6d, respectively).

**Figure 6:**
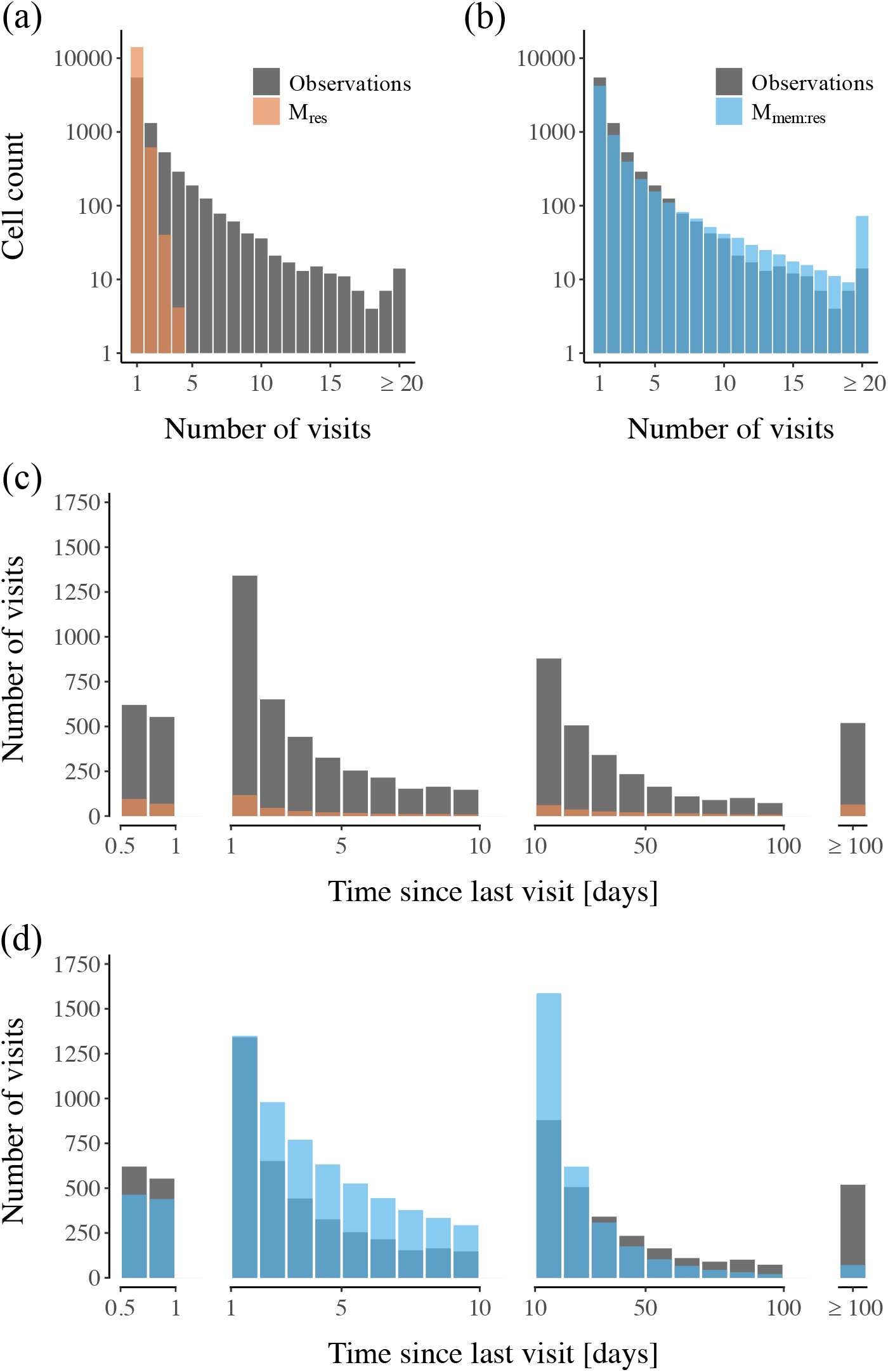
Emergent revisitation properties. The distributions of revisits (panels a and b) and time since last visit (panels c and d) are shown for observed roe deer movements (grey), the simulated trajectories from the resource-only model (M_res_; orange), and the simulated trajectories from the memory-based model (M_mem:res_; blue).

## Discussion

The past two decades have seen remarkable advances in the ability to monitor animal movements^30^, and the development of ever more complex and comprehensive mechanistic movement models. However, our understanding of the underlying biological determinants of home ranges – the most prevalent space-use pattern observed in animals – has been relatively limited^6,11,31^. In this study, we evaluated how memory-based movements can predict the formation of home ranges in nature by parametrizing a mechanistic movement model with empirical data from animals reintroduced into a novel environment. We found that an interplay between memory and resource preferences was the primary process influencing reintroduced roe deer movements (Fig. 2; H1), and that it led to the formation of characteristic home ranges, as observed in the released individuals (Figs 3 and 4; H2; see also Cagnacci *et al*. in prep). To our knowledge, this is the first demonstration that a mechanistic movement model parametrized with empirical movement data can capture patterns of home range formation in a non-territorial species.

We found that the emergent properties of the memory-based movement model, as opposed to a resource-only movement model, were realistic and similar to the patterns observed in the reintroduced roe deer (Figs 3–5). First and foremost, the memory-based simulations gave rise to spatially-restricted movements, as shown by the saturation of individual net squared displacement with time since release (Fig. 4). In addition, the model successfully reproduced the heterogeneity and complexity of observed movement patterns, including long-distance explorations, multiple areas of concentrated use, and patterns of revisitation (Figs 3 and 6).

Two approaches can be used to study the underlying determinants of animal space-use^32^: analyses inferring underlying movement parameters that capture observed space-use patterns i.e., pattern-oriented^3,8,31,33,34^, and analyses of individual movement trajectories to parametrise movement models^20,23,35^. In this study, we used the latter approach: characterizing the biological drivers of fine-scale behavioural decisions through the fitting of a mechanistic movement model to empirical trajectories, and subsequently evaluating resulting predictions of space-use properties. Although challenging, this approach is appealing because the space-use pattern itself is not fitted to data, but rather arises as an emergent property from the underlying movement process^32,35^.

Previous analyses have shown that memory influences the proximate behavioural decisions of free-ranging animals^19,20,22,23^. Our study extends these analyses in three major ways. First, we show that a movement model operating in a spatially continuous landscape not only accounts for the observed aggregate (population-level) patterns of space-use, but also yields realistic patterns of individual space-use (Figs 3 and 4). Second, our empirical setting of animals reintroduced into a novel environment allowed us to avoid the problematic issue of how to initialize memory-based movement models^19,20,23^ that has been invoked to explain the discrepancies between predicted and observed space-use patterns^22^. Third, as we discuss in more detail below, in addition to space-use patterns, the memory-based movement model captured several emergent characteristics of empirical roe deer trajectories (Figs 5 and 6). This provides confidence that the model’s realistic predictions of space-use are arising because the model closely approximates the key characteristics of individual movement behaviour that underlie the formation of the home ranges.

Patterns of animal space-use recorded by GPS-telemetry can be viewed as resulting from a sequence of movement decisions by the animal about how far to move, and in which direction i.e., sequences of movement distances and turning angles^9,36^. Our memory-based model was able to accurately characterize the distributions of both these quantities (Fig. 5). We found that a realistic, heavy-tailed distribution of step lengths emerged from the combination of a large, exponentially-weighted motion capacity (that accounts for rare, long steps), and memory-based attraction (that accounts for the high density of short steps). It has been suggested that characterizing step lengths with heavy-tailed Gamma or Weibull distributions could improve the predictive performance of empirically-parametrized movement models^20^. Here we show that accounting for memory makes this unnecessary. Furthermore, incorporating the effects of memory also gave rise to frequent reversals in movement directions (i.e., sharp turning angles) that closely matched the movement behaviour of released roe deer, even though the underlying redistribution kernel did not include any form of autocorrelation in movement directions.

Similarly, home ranges are thought to emerge from the revisitation of specific geographic locations (also referred to as movement recursions^29^), considered to be the visible manifestations of the influence of memory on movements^11,29^. Our results are consistent with this interpretation. The resource-only model led to very few revisits, while a revisitation behaviour similar to that observed in reintroduced roe deer emerged from the memory-based movement simulations (Fig. 6). The memory-based model predicted well the overall distribution of time since last visits, although it tended to underestimate both short (a day or less) and very long time since last visits (especially above 100 days) i.e., smaller variance that the observed pattern. The high density of short-term revisits in observed roe deer trajectories could be the result of daily, local movements such as an alternation between a foraging ground and a neighbouring area used for shelter^37^. The discrepancy observed for long-term revisits could instead be due to an artefact of the data collection (time since last visit can be overestimated in empirical data if visits occur between two successive GPS relocations), or could reflect biological factors that are not characterized in the model formulation (e.g., particular environmental conditions may be exploited through proximal mechanisms such as perception; see Avgar *et al*.^20^ for a cognitive model including perception and memory).

Similar to previous studies^15–17^, we hypothesized that home ranges emerge from the influence of a bi-component memory process in which reference memory captures the long-term attraction to previously-visited locations, while working memory accounts for a short-term repulsion (e.g., to adjust to resource dynamics). However, we found that it was not necessary to include a bi-component memory to give rise to home ranges. This result contrasts with the patch-to-patch transition model of Van Moorter *et al*.^15^, where absence of working memory leads to repeated utilization of a sole resource patch. In the memory-based movement formulation used here, an intrinsic component of resource preference ensures that all locations, including those that have not been visited, or that have been forgotten (i.e., whose reference memory is zero), have a non-zero probability of being visited at each time step.

When fitting the movement model to empirical data, reference memory was the most influential driver of roe deer movement (Table 1). The learning associated to the initial visit of any given location led to a 31.6-fold increase in its attraction (Fig. 2a), and hence to a substantial increase in its probability of being revisited in the future. We found that the influence of memory on movements was very strong despite the learning rate of reference memory being low. Although, learning was modelled as an exponentially saturating function of experience^38,39^, the low value of learning rate effectively meant that learning never approached its asymptote, and was essentially a quasi-linear function of number of visits. In this aspect, our findings provide support for the simple memory enhanced random walk formulation proposed by Tan *et al*.^40^. However, in contrast to Tan *et al*.^40^, our formulation also includes a spatial scale of learning parameter, which was strongly supported (Table 1), and implies that roe deer are likely to return not only to their previously visited locations but also to adjacent areas (Fig 2a).

Reference memory decayed relatively rapidly (half-life of 9.5 days) with time since last visit. This estimate is relatively consistent with the decay rate reported in a recent experimental study of roe deer foraging behaviour (half-life of 3.4 days)^21^, but contrasts markedly with the negligible decay of spatial memory over several months reported for bison (*Bison bison*)^19^ and woodland caribou (*Rangifer tarandus caribou*)^20^. Comparative studies may shed light on whether the factors underlying the differences in estimated memory decay rates are biological (e.g., variation in revisitation patterns linked to differences in movement rates and home range sizes), or methodological (e.g., between-model differences in the formulations of the cognitive processes).

Our results were, in contrast, much less sensitive to working memory. Working memory primarily influenced the duration that roe deer spent at visited locations (i.e., residence time). Because of its nearly instantaneous decay, the repulsion effect of working memory ceased as soon as the individual left a visited location. As a result, working memory did not influence the timing of revisits, which contrasts with predictions derived from theoretical movement simulations^15–17^. The probability that roe deer returned to previously visited locations decreased monotonically with time since their last visit (Fig. 5d), suggesting that a single memory component (reference memory, in our case) could capture roe deer revisitation patterns. Altogether, our results did not support the existence of characteristic multi-day revisitation periodicities, which would be expected if roe deer relied on working memory to optimally adjust their visits to underlying resource renewal dynamics. Because roe deer are very selective browsers, able to switch feeding between an important diversity of plant species^41,42^, it is indeed unlikely that their foraging behaviour is influenced by short-term dynamics of resource renewal. In contrast, a bi-component memory process may be more suited to model the foraging behaviour of species whose resources are concentrated within distinct, continuously-renewing patches such as grazing lawns in bison^43^ or geese^44^.

In our study, the estimated memory parameters gave rise to a strong attraction to familiar locations, consistent with published literature in roe deer^21,28^, and other ungulates^19,45^. Two main hypotheses have been formulated for the fitness benefits associated with site familiarity: (i) improved resource acquisition through the memorization of resource locations and attributes^10,15^, and (ii) predator avoidance through the knowledge of fine-scale variations in predation risk and of escape routes^46^. Previous work has shown that in roe deer, individuals rely on memory to efficiently track the spatio-temporal changes in food availability within their familiar environment^21^ but are also prone to elevated predation risk from Eurasian lynx (*Lynx lynx*) when outside of their familiar space^47^. These two benefits of site familiarity are difficult to disentangle in nature; in our study, both factors may likely have driven the revisitation patterns that contributed to the emergence of roe deer home ranges.

Our analysis also revealed the resource preferences of roe deer in our study area. First, roe deer exhibited strong preference for intermediate slope steepness (Fig. 2b). Their avoidance of flat areas is likely explained by the fact that, in the rugged landscape of Aspromonte National Park, anthropogenic disturbances such as roads and logging activities^48^ were concentrated along valley bottoms, as well as high plateaus. In other ecological systems, these topographic features have also been associated with elevated predation risk from wolves^49^. Their avoidance of steep slopes is consistent with roe deer natural history (long limbs, and short and narrow hoofs not adapted to climbing), and its unsuitability supported by the occurrence of two mortality cases linked to falls during the reintroduction project (*S. Nicoloso pers. comm*.). Second, roe deer preferred areas of intermediate-to-high tree cover (Fig. 2c), a finding that is consistent with published literature on roe deer resource selection^50–52^. Intermediate cover values may indicate heterogenous environments rich in ecotones, which provide abundant browsing resources^53^. Third, we found that roe deer strongly preferred reforested areas with young deciduous trees (Fig. 2d), which is likely because these areas provide both cover and abundant browse^42^. Fourth, roe deer avoided agricultural areas (and associated pastures and settlements; Fig. 2d) in agreement with existing literature^50,51^.

Despite qualitative similarities between the resource-only and memory-based model formulations, the effect sizes of resource preference parameters were consistently smaller for the memory-based model than for the resource-only model (Fig. 1; Fig. 2b-d). In absence of memory, the relative attraction (and hence probability of use) of equally-distant locations solely depends on their respective resource attributes. In contrast, when memory processes operate the relative attraction is partitioned between two interacting components: resource attributes (i.e., resource effect), and memory (i.e., site familiarity effect) – thereby reducing the influence of resources *per se*. Because animal home ranges ultimately emerge as the revisitation of familiar, beneficial resources^10,15^, disentangling the influence of resources from that of site familiarity is challenging in nature. In particular, where important resource drivers are omitted – either because they are unknown or because they are not measured – the attraction for familiar areas can be confounded with attraction for unaccounted resources (i.e., a spurious familiarity effect^54^). Further progress to characterize the interplay between memory and resource preferences will be contingent on the ability to identify and quantify underlying spatio-temporal variation in resource patterns. In this context, combining mechanistic movement models with *in situ* experimental resource manipulations appears a promising way to disentangle the effects of memory from the effects of resources^21,28^.

Connecting animal movement behaviour to space-use patterns and, ultimately, population dynamics is a long-term challenge that promises to provide a unifying theory for animal ecology^55^. In this study, we demonstrated that the interplay between memory and resource preferences is sufficient to explain the formation of animal home ranges following reintroduction to a novel environment, and thus contributing to our understanding of the space-use implications of movement behaviour. The approach utilised here could be expanded to model the interconnections between movement behaviour and energy acquisition and consumption^56^, providing a framework to quantitatively characterize the fitness, and demographic consequences of animal movement patterns, and space-use^57^.

## Methods

### Roe deer reintroduction

After being extirpated in most of its southern distribution range during the 19^th^ century, a roe deer reintroduction project was undertaken by the Aspromonte National Park (AspNP; Calabria, Italy; Supplementary S2: Fig. S1) between 2008 and 2011. Ninety-two roe deer were captured in Sienna County (Tuscany, Italy), of which seventy-five were hard-released at four sites in the south-west portion of the AspNP (47 females and 28 males). The remaining seventeen either died during translocation or were not genotyped as *Capreolus capreolus italicus*, the roe deer subspecies native to the Italian peninsula.

The AspNP is 640 sq.km and is characterized by the rugged Aspromonte mountain range peaking at 1955 m a.s.l, and alternating gorges and torrent river valleys. The climate is Mediterranean with precipitations concentrated in winter, leading to irregular snow cover above 1000 m a.s.l, and dry and warm summers (annual precipitation: 826 mm; temperature: −0.8/5.4°C in January, 14.9/23.0°C in August; Gambarie, 1300 m a.s.l). The significant topography within the region gives rise to a diverse vegetation cover^58^ ranging from temperate mountain forests (e.g., European beech *Fagus sylvatica*, silver fir *Abies alba* and alder *Alnus sp*.) to dry pine and oak forests (e.g., Calabrese black pine *Pinus laricio*, Mediterranean oaks *Quercus ilex* and *Quercus suber*), and high Mediterranean maquis (e.g., strawberry tree *Arbutus unedo*, heather *Erica arborea* and myrtle thickets *Myrto-Pistacietum lentisci*). The region also includes small-scale, mixed agriculture, orchards and plantations (e.g., chestnut *Castanea sativa*, and olive groves), pastures at high elevation, as well as small settlements at the margins of the park. Wild boar (*Sus scrofa*) is the dominant wild ungulate in the study area (red deer, *Cervus elaphus* are locally extinct). Wolves (*Canis lupus*) are the only natural predators of adult roe deer, although red fox (*Vulpes vulpes*) may predate upon fawns. Hunting is forbidden within the national park.

### Empirical data

The movements of twenty-seven individual roe deer were monitored after their release via telemetry. Roe deer were fitted with GPS-GSM collars scheduled to acquire one relocation every 30 min during the first month after release, and at six-hour intervals thereafter (schedule: 00:00, 06:00, 12:00, 18:00 UTC). For the purpose of our analysis, we retained all animals for which we could obtain a trajectory of at least 30 days with a high acquisition success rate (> 85%). This choice led to the exclusion of ten individuals – seven died in the first month after release and three had malfunctioning collars. Our final sample consisted of 17 roe deer (15 adults: 11 females, 4 males; 2 subadult males), tracked for an average of 281.82 days (*σ* = 167.37, minimum = 39, maximum = 624; Supplementary S3: Table S1).

We regularized the trajectories to a homogeneous relocation interval of six hour and did not interpolate the missing relocations. The final dataset consisted of 19,186 GPS relocations (acquisition success rate = 93.61%). Roe deer step length averaged 140.04 m between two successive relocations (*σ* = 267.37, maximum = 6254.68 m).

We analysed the movement behaviour of the reintroduced roe deer within a rectangular area (40.8 x 30 km; 1,224 sq.km; Supplementary S2: Fig. S1), that encompassed all available roe deer GPS locations and a buffer of 7 km (more than the longest observed step length). Given the average movement distance of the reintroduced roe deer, and the high landscape heterogeneity of our study area, the landscape was represented at a spatial resolution of 25 x 25 m.

The resource preference component of the mechanistic movement model included both topographic (slope) and landcover variables (tree cover, agriculture and reforested landcover). We selected these variables a priori as they are known predictors of roe deer movement and resource selection^42,48,51,52^, and a preliminary step selection analysis (SSA^59^; results not shown) ascertained their relevance in our study system.

We obtained the slope layer from the European Union Digital Elevation Model EU-DEM v1.0^60^. Slope ranges from 0 to 90°, and was available at a 25 m spatial resolution. We obtained an estimate of tree cover from Copernicus pan-European, high-resolution layers^61^, 2012 reference year. Tree cover ranges from 0 to 100%, and was resampled at 25 m from a native spatial resolution of 20 m. Following preliminary SSA explorations, we (1) calculated tree cover at a grain of 325 m (i.e., each squared cell of 25 m averaged tree cover within a larger 325 x 325 m area) and (2) included in the model both linear and quadratic terms for slope and tree cover.

Two landcover data sources were available for our study area – a botanical map of high biological detail and fine spatial resolution (94 categories; 0.05 ha mapping unit) for the Aspromonte National Park^58^, and the coarser CORINE landcover classification (45 categories; 25 ha mapping unit) for the entire study area^62^, 2012 reference year. Preliminary SSA conducted within the park boundaries suggested that roe deer selected for areas reforested with deciduous trees (*Alnus cordata, Juglans regia* and *Prunus avium*; hereafter referenced to as *Reforested*), and avoided spatially-dominant agriculture areas (olive groves, cultivated fields, mixed agriculture), and more localized pastures and anthropized areas (hereafter referenced to as *Agriculture*). Outside the park, we assumed that there were no *Reforested* areas, and used CORINE to map *Agriculture* – choices that we validated by visually inspecting the satellite images in the vicinity (< 1 km) of the roe deer relocations outside of the park.

### Modelling approach

We modelled the movement of reintroduced roe deer using an individual-based, spatially explicit *redistribution kernel* combining spatial memory and resource preferences. Specifically, we defined the probability of moving between the relocation **x**_*t*−1_ and the relocation **x**_*t*_ (as it is standard: **x** = (*x, y*)), as the normalized product of an *information-independent movement kernel*^27^, *k*(**x**_*t*_; **x**_*t*−1_, *θ*_1_), and a *cognitive weighting function, w*(**x**_*t*_; *t*, *θ*_2_)^20,26^:

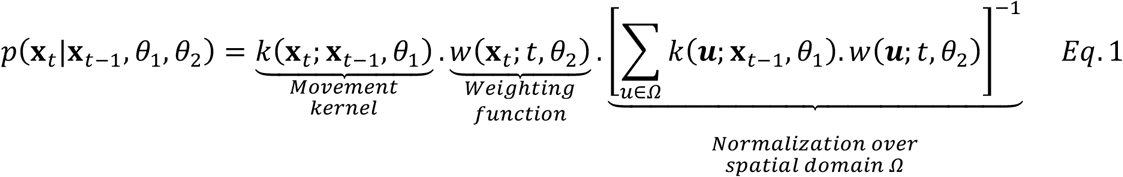

with ***u*** = (*x, y*) denoting all the locations within the within the spatial domain *Ω*, and *θ*_1_ and *θ*_2_ the ensemble of parameters governing the movement kernel, and the weighting function, respectively.

### Motion capacity – the information-independent movement kernel

The information-independent movement kernel characterizes the movement of an animal independently of its cognitive abilities and of the surrounding landscape, and therefore quantifies its motion capacity^20^. It is obtained through the product of two probability distributions: step length, *S*, and movement direction, Φ. Here, we modelled roe deer step length using a truncated Weibull distribution. The Weibull distribution is governed by two parameters – the shape (*κ_S_* > 0) and the rate (*λ_S_* ≥ 0), and can account for both a high density of short movements and rare, long movements (i.e., heavy tail), typical of empirical data of animal movement^63^. To reduce the computational power required for model fitting, we assumed that roe deer movement probability was zero beyond 7 km (maximum observed step length = 6.25 km). The resulting step length distribution for any location ***u*** is given by:

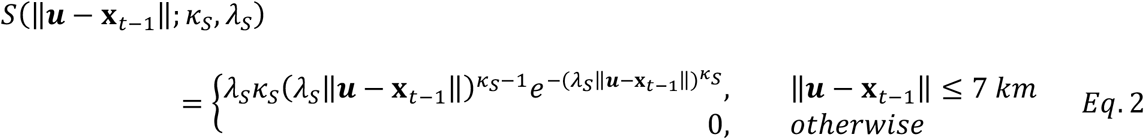

We modelled roe deer movement directions as a circular normal distribution such that:

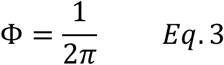

It follows that the information-independent movement kernel is given by:

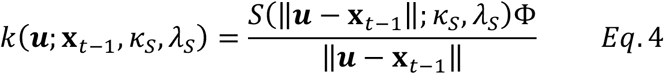

where the denominator ||***u*** – **x**_*t*−1_|| translates polar coordinates into Euclidean coordinates i.e., the conversion of a probability of moving a given distance and direction to a probability of moving to a particular area^9^. Given the temporal resolution of our movement data (every 6 h), we ignored serial correlation in movement direction. Because we fitted our mechanistic movement model to observed movement data in a discretized landscape (square cells of resolution 25 m), we transformed the GPS relocations in the continuous space, **x**_*t*_, to the centroid of the overlapping cell (see Supplementary S4 for the correction required to calculate the movement kernel on the location currently used by the animal).

### Interplay between memory and resource preferences – the cognitive weighting function

The interaction between the landscape and the animal cognitive abilities was represented via the weighting function *w*. We assumed that animal movement was influenced by memory, *m*(***u***; *t*), and that, in absence of such information, animals may visit locations in proportion to their intrinsic resource preference value:

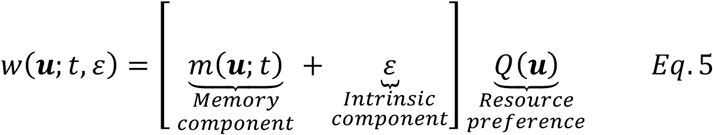

with *ε*, the intrinsic component of resource preference. In our model formulation, it is not the absolute value of memory that defines its influence on movement, but rather its value relative to the intrinsic component of resource preference i.e., scaled to the attraction of similar resource conditions in absence of memory. We modelled the preference for location *u, Q*(***u***), using an exponential resource selection^64^:

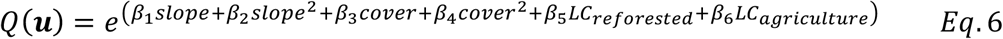

with *β_Y_* the selection coefficient for resource variable *i* – slope (linear and quadratic terms), tree cover (linear and quadratic terms), and reforested and agriculture landcovers – evaluated at location ***u***.

We modelled memory, *m_R_*(***u***; *t*), as a bi-component mechanism^15–17^. Reference memory, *m_R_*(***u***; *t*), is the long-term memory of previously-visited spatial locations and has an attractive effect. By contrast, the working memory, *m_W_*(***u***; *t*), encodes the short-term, temporary repulsion of previously visited locations. The combined memory map is given by:

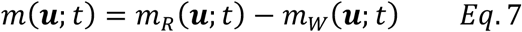

The dynamics of both memory components are governed by learning (i.e., acquisition of information) and decay or forgetting (i.e., loss of information). The learning curve was represented by an asymptotically increasing function of experience^38,39^. Specifically, we formulated learning as an exponentially saturating process with an associated spatial scale such that animals experience maximum learning at their current position, but also gain information about surrounding areas. Decay was modelled as a negative exponential of time since last visit^65,66^. Together this yields the following equations for the dynamics of memory across space ***u***, given the animal’s current position **x_*t*_**:

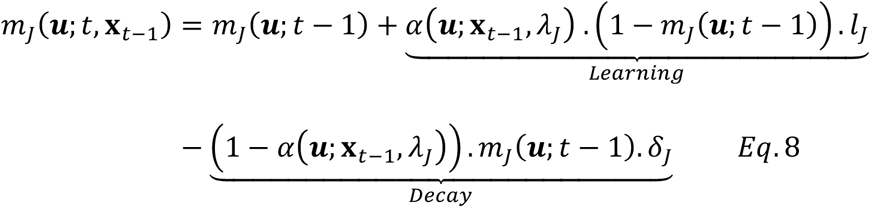

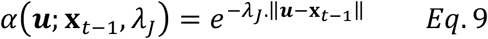

where *J* = *R* and *J* = *W* for reference and working memory, respectively; *l_R_* and *l_W_* are the rates of learning for reference and working memory; the functions *α*(***u***; **x**_*t*−1_, *λ_R_*) and *α*(***u***; **x**_*t*−1_, *λ_W_*) respectively describe how the rates of reference and working memory acquisition attenuate as a function of distance from the animal’s previous position (modelled via negative exponential functions); and *δ_R_* and *δ_W_* determine the rates at which the two forms of memory decay over time. According to their biological definition^15,16^, reference memory is always larger than working memory, thus imposing the following constraints: *l_R_* ≥ *l_W_*, *δ_R_* ≤ *δ_W_* and *λ_R_* ≤ *λ_W_*. For missing relocations, no learning occurred but memory decay took place.

### Model fitting

We fitted two models representing competing hypotheses pertaining to the biological processes influencing the movements of reintroduced roe deer: resource-only (M_res_), and interplay between memory and resources (M_mem:res_). For the M_res_ model, the memory parameters were omitted (i.e., no memory learning; *l_R_* = *l_W_* = 0). We estimated the model parameters through maximum-likelihood inference. The likelihood function for the parameter set of the information-independent movement kernel *θ*_1_ = *κ_S_, λ_S_*, and of the cognitive weighting function, *θ*_2_ = (*l_R_, l_W_, δ_R_, δ_W_, λ_R_, λ_W_, ε, β*_1_: *β*_6_), is given as:

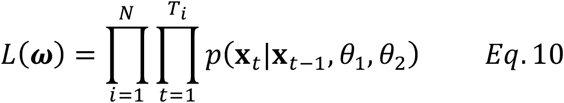

with *N* the number of animals (i.e., 17) and *T_i_* the number of relocations for animal *i*. Missing GPS relocations were omitted from the likelihood function. We estimated the global minima of the log-likelihood function [*logL*(***ω***); i.e., the objective function] using the particle swarm optimization algorithm (PSO^67^; see Supplementary S5 for details). We calculated 95% marginal confidence intervals (CIs) via an asymptotic normal approximation of the objective function in the neighbourhood of the global minima. We then evaluated the contribution of each variable to the model support by calculating the delta Akaike Information Criterion^68^ of the reduced model (i.e., excluding the variable of interest) relative to the full model.

### Movement simulations and emergent properties

We evaluated whether the two parametrized movement models (M_res_ and M_mem:res_) could characterize the spatial behaviour of reintroduced roe deer by means of movement simulations. For each monitored roe deer, we ran 30 movement simulations (17 animals x 30 runs = 510 simulated trajectories per model), initiated on the first observed GPS relocation of each individual (i.e., in the vicinity of the release site). At each time step, a spatial location was randomly selected according to the probabilities defined by the parametrized redistribution kernel. Simulations ran for a duration equivalent to that of the observed roe deer trajectories.

We compared observed and simulated trajectories using key emerging properties. First, to evaluate the emergence of spatially-restricted movements, we compared the temporal trend in net squared displacement (NSD). NSD was calculated as the squared distance between the individual position at time *t*, **x**_*t*_, and the trajectory start position, **x**_0_:

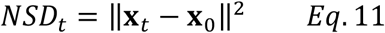

At the population-level, we computed the mean NSD for the 17 released roe deer as a 5-day running mean to remove individual noise. For the simulations, we calculated the 5% and 95% confidence bounds for the mean NSD via bootstrapping (1000 random samples of 17 simulated trajectories). Second, we evaluated whether the parametrized movement models captured the empirical distributions of emergent movement properties. We calculated step length as the Euclidean distance between two successive relocations, and turning angle as the angle in radians between the directions of two successive steps (ranges from −*π* to *π*; 0 indicating no directional change). Third, we investigated whether the parametrized movement models captured roe deer revisitation behaviour (i.e., movement recursions^29^). Revisits, defined as returns to a previously visited area, occurred when an animal (observed or simulated) used a 25 x 25 m spatial cell that had been last visited within > 6 hours (i.e., temporally-disjointed use of a specific location). For each visited cell along the trajectory, we computed its total number of revisits (0 indicating a single visit) and their associated time since last visit.

Equations 1–10 were solved numerically, and simulations performed in C++. The parameters were estimated using the PSO algorithm, implemented within the Global Optimization Toolbox, MATLAB R2017b (MathWorks, Natick, Massachusetts, USA). The optimization ran on a computer cluster using the Distributed Computer Server^69^. We calculated the CIs, produced the effect size plots and comparison between observed and simulated trajectories in R^70^.

## Acknowledgments

N. Ranc was supported by a Harvard University Graduate Fellowship and a Fondazione Edmund Mach International Doctoral Programme Fellowship. F. Cagnacci was supported by the Sarah and Daniel Hrdy Fellowship 2015-2016 at Harvard University OEB during part of the development of this manuscript. We thank the Aspromonte National Park (Calabria, Italia) for promoting and financially supporting the roe deer reintroduction project, and the applied ecology (agriculture, forestry and wildlife management) cooperative *D.R.E.Am. Italia* (Tuscany, Italy). We especially and warmly thank Sandro Nicoloso and Lillia Orlandi (*D.R.E.Am. Italia*), and Antonino Morabito (*Legambiente*). We are also very grateful to P. Krastev (Harvard Research Computing), A. La Fata, B. Ölveczky, N. Pierce and J.W. Cain for their valuable suggestions.

## Data Availability Statement

The datasets generated and/or analysed during the current study are available from the authors on request.

## Ethical Statement

The roe deer (*Capreolus capreolus spp. italicus*) reintroduction project was carried out under the technical approval and ethical requirements of the Institute for Environmental Protection and Research (ISPRA, public body for research under the vigilance of the Italian Ministry of the Environment; protocols #47427 and #45826). The capture of roe deer in the population of origin were conducted under authorization #1555 by the Director of the Wildlife Resources and Natural Reserve Service of the Province of Sienna.

