## Supplementary Materials for "Memory drives the formation of animal home ranges: evidence from a reintroduction"

**Supplementary S1: Movement trajectories**

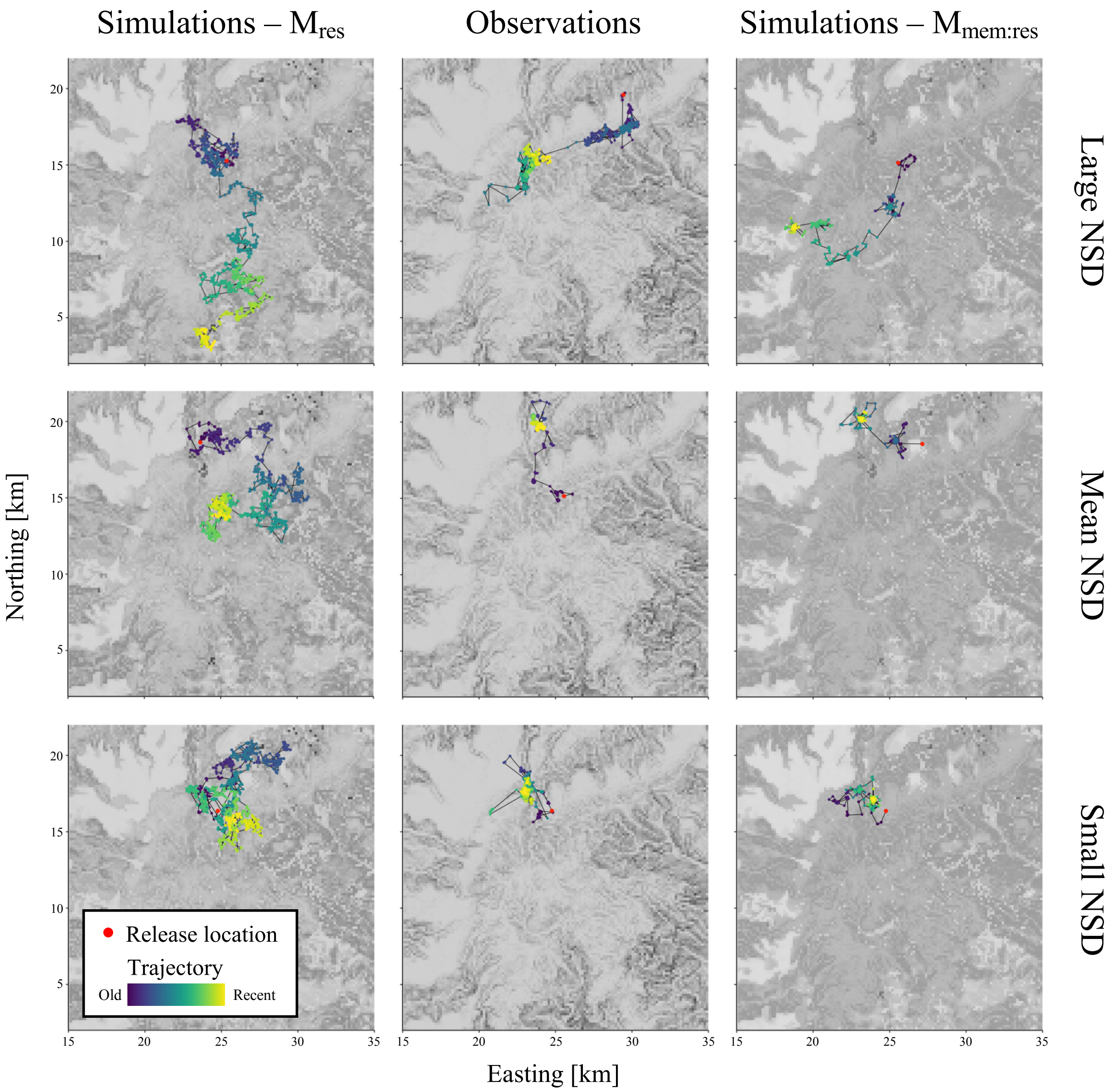


**Figure S1: Movement trajectories**. The trajectories for the resource-only simulations ($M_{res}$; left column), observed roe deer movements (central column), and memory-based simulations ($M_{mem:res}$; right column) are shown for three categories of final net squared displacement – final location far from the release location (top row), at average distance (central row) or close (bottom row). The release location is shown as a red dot and the time since release illustrated as a colour gradient (blue = old, yellow = recent). The trajectories were selected from the samples displayed on Figure 4 (see Main text).

**Supplementary S2: Study area map**

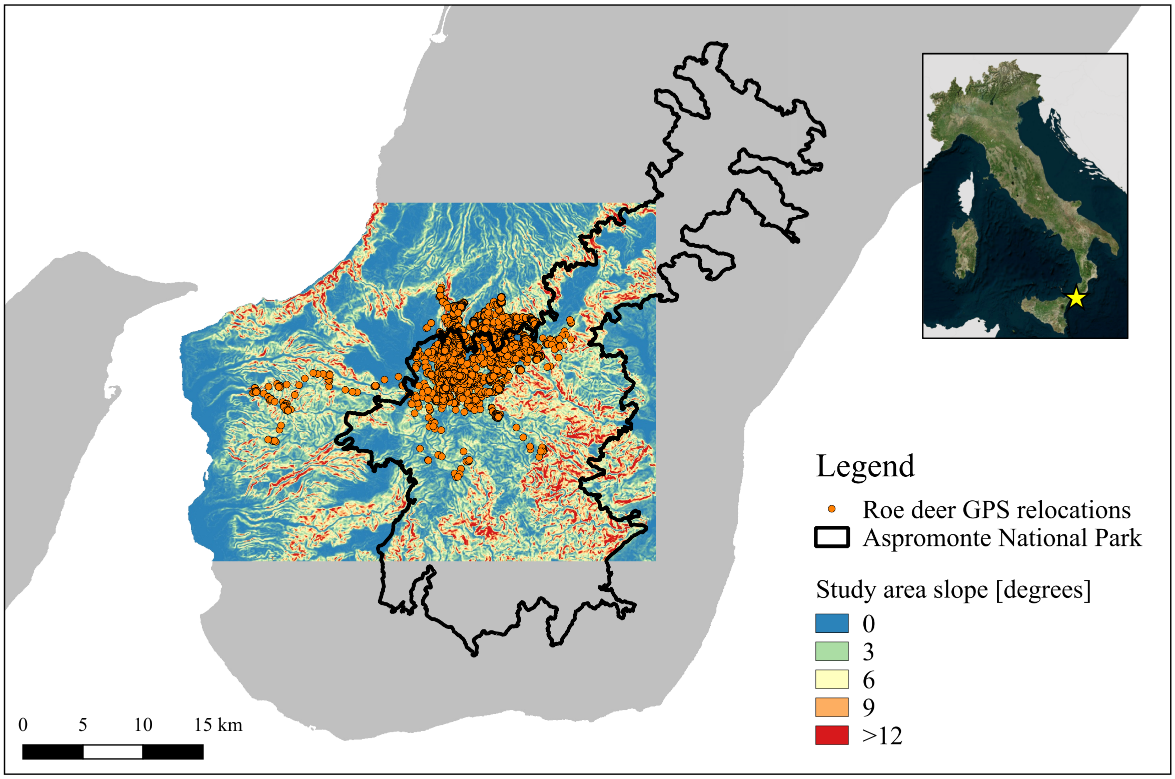


**Figure S1: Map of the study area.** The study area is located in Calabria region (Southern Italy) and covers the Southwestern part of the Aspromonte National Park (thick black line). Slope is shown in coloured gradient, and roe deer GPS locations are shown as orange dots.

### Supplementary S3: Roe deer movement data

**Table S1: Data summary for the 17 roe deer included in the study**.

| Individual | Sex | Age | First location | Days of monitoring | Relocation success rate |
|---|---|---|---|---|---|
| 1182 | f | adult | 16/11/2008 12:00 | 351 | 0.98 |
| 1183 | f | adult | 16/11/2008 12:00 | 362 | 0.98 |
| 1196 | m | adult | 16/11/2008 12:00 | 105 | 0.96 |
| 1199 | m | subadult | 16/11/2008 12:00 | 624 | 0.87 |
| 1181 | f | adult | 02/03/2009 12:00 | 577 | 0.90 |
| 1187 | f | adult | 02/03/2009 12:00 | 181 | 0.94 |
| 1195 | f | adult | 20/03/2009 12:00 | 238 | 0.90 |
| 1200 | m | subadult | 20/03/2009 12:00 | 182 | 0.93 |
| 1186 | f | adult | 10/02/2010 12:00 | 363 | 0.95 |
| 1193 | f | adult | 10/02/2010 12:00 | 363 | 0.94 |
| 1194 | f | adult | 12/02/2010 18:00 | 361 | 0.98 |
| 1197 | m | adult | 10/02/2010 12:00 | 363 | 0.93 |
| 1189 | f | adult | 27/02/2010 12:00 | 154 | 0.94 |
| 1190 | f | adult | 27/02/2010 12:00 | 346 | 0.96 |
| 1458 | m | adult | 16/02/2011 12:00 | 67 | 1.00 |
| 1459 | m | adult | 16/02/2011 12:00 | 39 | 0.99 |
| 1460 | f | adult | 19/11/2011 12:00 | 115 | 0.90 |

Dashed lines separate the different release events. Roe deer 1194 was released in the same time as roe deer 1186, 1193 and 1197 but the GPS collar malfunctioned during the first two days.

**Supplementary S4: Correction factor for the information-independent movement kernel**

When evaluating the information-independent movement kernel for the spatial cell currently used by the animal (i.e., when $\boldsymbol{u} = \mathbf{x}_{t-1}$), we set the distance $\|\boldsymbol{u} - \mathbf{x}_{t-1}\|$ to $\vartheta$, a correction factor equal to the mean distance moved in continuous space when an animal remains in the same pixel in the discretized landscape i.e., $\boldsymbol{u} = \mathbf{x}_{t-1}$. In a cartesian landscape $(x, y)$ of resolution $r$ (here, $r = 25$ m), $\vartheta$ is given by:

$$\vartheta = \left[ \int_{x=-\frac{r}{2}}^{\frac{r}{2}} \int_{y=-\frac{r}{2}}^{\frac{r}{2}} \frac{S(Dist; \kappa_S, \lambda_S)\Phi}{Dist} Dist.dxdy \right] \times \left[ \int_{x=-\frac{r}{2}}^{\frac{r}{2}} \int_{y=-\frac{r}{2}}^{\frac{r}{2}} \frac{S(Dist; \kappa_S, \lambda_S)\Phi}{Dist} dxdy \right]^{-1} \qquad Eq.1$$

with $Dist = \sqrt{x^2 + y^2}$. Since $\Phi = \frac{1}{2\pi}$ i.e., a constant, the above equation reduces to:

$$\vartheta = \left[ \int_{x=-\frac{r}{2}}^{\frac{r}{2}} \int_{y=-\frac{r}{2}}^{\frac{r}{2}} S(Dist; \kappa_S, \lambda_S) dxdy \right] \times \left[ \int_{x=-\frac{r}{2}}^{\frac{r}{2}} \int_{y=-\frac{r}{2}}^{\frac{r}{2}} \frac{S(Dist; \kappa_S, \lambda_S)}{Dist} dxdy \right]^{-1} \qquad Eq.2$$

The inclusion of the $\vartheta$ correction factor significantly improved the match between theoretical and simulated step length distributions (compare Figs S1a and S1b).

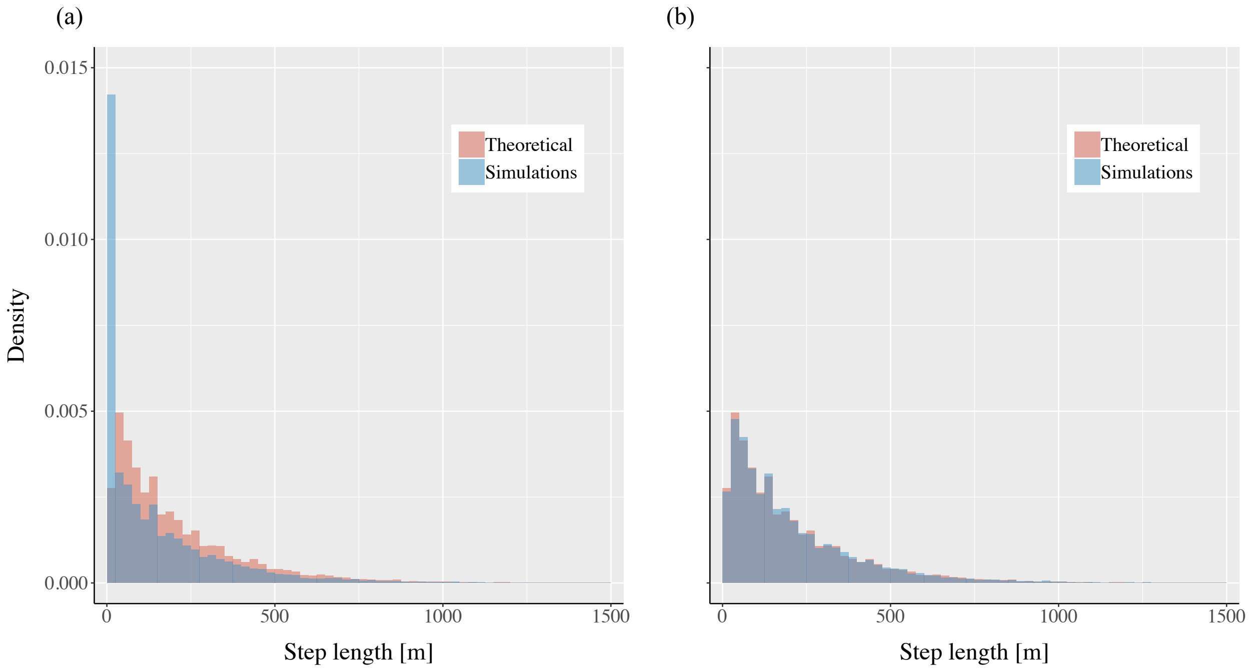


**Figure S1: Visualization of the effects of the correcting factor.** We compare a theoretical, two-dimensional Weibull step length distribution (shape $\kappa_S = 1$ and rate $\lambda_S = 0.005$; in red) with movement simulations (blue), generated using either an uncorrected (panel a) or corrected (panel b) movement kernel.

**Supplementary S5: Particle swarm optimization**

We used a non-linear, heuristic article algorithm, particle swarm optimization (PSO) to estimate the best set of parameters of our movement models. PSO relies on collective behaviour to converge to a solution into the parameter search-space (see Poli et al.[1] for a review). Because our problem contains many dimensions (15), we initially chose a large swarm (150 particles) for a duration of 400 iterations, which proved sufficient for our task. We let the PSO reach the maximum number of iterations, although convergence to two significant digits was usually achieved after 200-250 iterations. We dampened the velocity of the particles in the search space to stabilize the algorithm i.e., a constriction PSO[1], and used the optimal settings described by Clerc & Kennedy[2] listed below:

Constant, non-adaptive inertia ($\omega$) = 0.7298

Self-adjustment weight ($\emptyset_1$) = 1.49618

Social adjustment weight ($\emptyset_2$) = 1.49618

We used the algorithm available in the MATLAB Global Optimization Toolbox (MathWorks, Natick, Massachusetts, USA) and set the neighbourhood fraction to the default value of 0.25. To facilitate and increase the speed of the optimization, we constrained the parameter search-space on the most complex model as,

$$\kappa_S \in [0.3,3]$$

$$\lambda_S \in [0.0001,0.1]$$

$$\beta_1 : \beta_6 \in [-3,3]$$

$$l_R, l_W \in [0,1] \text{ with } l_R \geq l_W$$

$$\delta_R, \delta_W \in [0,0.999] \text{ with } \delta_W \geq \delta_R$$

$$\delta_R, \delta_W \in [0,1] \text{ with } \lambda_W \geq \lambda_R$$

$$\gamma \in [0,1]$$
